# Apoptotic factors and mitochondrial complexes assist determination of strain-specific susceptibility of mice to Parkinsonian neurotoxin MPTP

**DOI:** 10.1101/2022.10.05.511004

**Authors:** Haorei Yarreiphang, D J Vidyadhara, Anand Krishnan Nambisan, Trichur R Raju, BK Chandrashekar Sagar, Phalguni Anand Alladi

**Author notes:** Equal 1^st^ authors (Haorei Yarreiphang, D J Vidyadhara). **Corresponding Author:** Dr. Phalguni Anand Alladi, Senior Scientific Officer-Scientist ‘F’, Department of Clinical Psychopharmacology and Neurotoxicology. Formerly at- Department of Neurophysiology, National Institute of Mental Health and Neuro Sciences, Hosur Road Bangalore, India (Phalguni Anand Alladi). **HY:** Zoology Department, Hansraj College, University of Delhi 110007, Delhi **VDJ:** Departments of Neurology and Neuroscience, Yale University School of Medicine, New Haven, Connecticut, USA.

## Abstract

Identification of genetic mutations in Parkinson’s disease (PD) promulgates the genetic nature of disease susceptibility. Resilience-associated genes being unknown till date, the normal genetic makeup of an individual may be determinative too. Our earlier studies comparing the substantia nigra (SN) and striatum of C57BL/6J, CD-1 mice and their F1-crossbreds demonstrated neuroprotective role of admixing, against the neurotoxin MPTP. Further, the differences in levels of mitochondrial fission/fusion proteins in the SN of parent strains, imply effect on mitochondrial biogenesis. Our present investigations suggest that the baseline levels of apoptotic factors Bcl-2, Bax and AIF differ across the three strains and, are differentially altered in SN following MPTP-administration. The reduction in complex-I levels exclusively in MPTP-injected C57BL/6J, reiterate mitochondrial involvement in PD pathogenesis. The MPTP-induced increase in complex-IV, in the nigra of both parent strains may be compensatory in nature. Ultrastructural evaluation showed fairly preserved mitochondria in the dopaminergic neurons of CD-1 and F1-crossbreds. However, in CD-1, the endoplasmic reticulum demonstrated distinct luminal enlargement, bordering onto ballooning, suggesting proteinopathy as a possible initial trigger.

The increase in α-synuclein in the pars reticulata of crossbreds suggests a supportive role for this output nucleus in compensating for the lost function of pars compacta. Alternatively, since α-synuclein over-expression occurs in different brain regions in PD, the α-synuclein increase here may suggest similar pathogenic outcome. Further understanding is required to resolve this biological contraption. Nevertheless, admixing reduces the risk to MPTP by favouring anti-apoptotic consequences. Similar neuroprotection may be envisaged in the admixed populace of Anglo-Indians.

## Introduction

Higher prevalence of Parkinson’s disease (PD) in the Caucasian populations has been a cause for concern for the health care policymakers in developed countries. Although there is relatively lower prevalence in Asians (Abbas et al., 2017) compared to Caucasians (von Campenhausen et al., 2005; Balestrino and Schapira, 2020), Asian-Indians (Das et al., 2010; Gourie-Devi M., 2014) and other non-white populations, it is predicted that the total number of patients are likely to increase in the next two decades in the highly populous Asian countries. It is hence pertinent to understand the factors that discriminate the vulnerable and resistant populations, to design necessary policies. PD related GWAS studies in European, Japanese and Han Chinese populations propose a strong involvement of proteins like α-synuclein, LRRK-2 and MCCC1 in the disease progression (Foo et al., 2017). On the contrary, there are few studies that explain the molecular mechanisms of resistance to PD.

Loss of mitochondrial function is implicated in PD pathogenesis (Banerjee et al., 2009b) confirmed by the link between pesticide use and increased risk of PD, as well as complex I dysfunction (Schuh et al., 2009). The primary evidence of mitochondrial abnormalities in PD was provided while modelling MPTP-induced Parkinsonism, which mainly targets complex-I of the electron transport chain (ETC) (Langston et al., 1983). Landmark studies revealed depletion in the complex-I expression in the SNpc of the autopsied brains of PD patients (Schapira et al., 1990a; 1990c; review by Toulorge et al., 2016). Mitochondria, harbor low levels of α-synuclein, the latter upon aggregation cause mitochondrial complex-I deficits and oxidative stress (Parihar et al., 2008). Several studies over decades, have shown that such dysfunction results in mitochondrial dysmorphogenesis, as seen in various experimental (Tanaka et al., 1988) and patient tissues (Hatton et al., 2020).

Differences in mitochondrial parameters may influence the apoptosis index of the neurons. Apoptosis may also occur in caspase-independent manner, i.e. via the release of apoptosis-inducing factor (AIF) from mitochondria, resulting in neurodegeneration (Bano & Prehn, 2018; Boujrad et al., 2007). AIF is a mitochondrial oxidoreductase that contributes to the assembly of the respiratory chain and apoptosis. Its deficiency leads to severe mitochondrial dysfunction, resulting in neurodegeneration in model organisms as well as in humans. The neuroprotective ability of the mitochondrial protein Parkin is contingent upon its cooperation with AIF (Guida et al., 2019).

Amongst the apoptosis related factors, Bcl-2 balances autophagy (Liu et al., 2018) and has therapeutic implications in PD (Czubowicz et al., 2019). As per a classical study, higher Bcl-2 level in PD brain was a compensatory neuroprotective mechanism (Jenner and Olanow, 1996). Regulation of Bcl-2, Bax and cytochrome-C through gene expression substantially enhanced motor functions and restored antioxidant enzyme activity in chronic MPTP-mice model of PD (Rekha and Selvakumar, 2014). Alteration of JNK/SAPK and ratio of Bcl-2 to Bax caused dopamine-induced apoptosis and continues to be an important parameter in drug discovery (Bekker et al., 2021).

The protein α-synuclein is involved in neurotransmitter release, acts as an uptake modifier and participates in vesicle trafficking and refilling in association with SNARE-complexes at the synapses. Although present in a monomeric form in normal physiological conditions, it has a tendency to aggregate (Irwin et al., 2013) and is a major player in neuronal death in PD (Irwin et al., 2013; Michel et al., 2016; Spillantini et al., 1997). Misfolding of α-synuclein induces cytotoxicity and affects all routine cellular processes. Sun et al. (2019) induced mutations to disrupt the interaction between C-terminus of α-synuclein with VAMP2 and implicated the latter in the inhibition of synaptic vesicle exocytosis by over-expressed synuclein. In a recent study on a drosophila model, α-synuclein overexpression caused mitochondrial fragmentation through the actin cytoskeleton (Ordonez et al. 2018). Yet, the role of α-synuclein in differential susceptibility is not completely understood.

There is ever expanding evidence of a functional association between PD and the gut microbiome opening up a possibility that the latter may potentially modulate the onset and or exacerbation of the disease and hence contribute to the susceptibility index of an individual. One of the essences of microbiota modulating the host responses is through the secretion of short chain fatty acids (Unger et al., 2018). Since diet and diurnal variations can be better controlled in an animal model, it would be ideal to use them to explore if any differences exist between different mice strains in terms of their gut microbiome.

In the present study we examined the effects of MPTP on apoptotic properties by assessing the apoptosis related protein AIF; pro and anti-apoptotic proteins Bcl-2 and Bax in the nigral dopaminergic neurons by immunofluorescence, confocal microscopy and western blotting. We further estimated the α-synuclein expression in SNpc and SNpr by semi-quantitative confocal microscopy. We assessed the complex I and IV activities followed by the ultra-structural alterations in the nigral DA neurons by transmission electron microscopy. We analyzed the sterile fecal pellets for the gut microbiome. The comparisons were made in C57BL6J and CD-1 mice strains, which exhibit differential susceptibility to MPTP; in addition to their F1 (first generation) crossbred progeny in response to MPTP.

## Materials and Methods

### Mice strains

The parent strains (C57BL/6J and CD-1) as well as the F1 progeny i.e. F1X1 and F1X2 (Vidyadhara et al., 2017; Vidyadhara et al., 2019) were housed in standard mice cages with ad-libitum access to food and water. All the experiments were carried out during the light period (08:00-18:00h) in accordance to the guidelines of the Committee for the Purpose of Control and Supervision of Experiments on Animals (CPCSEA), New Delhi, India, which adhere to the guidelines of National Institute of Health, USA.

### Immunofluorescence labelling and quantification

Adult mice [15-17 weeks, n=6/strain/experimental condition (C57BL/6J; CD-1; F1X1 and F1X2; control and MPTP injected)] following halothane inhalation were perfused intracardially with saline followed by 4% buffered paraformaldehyde (0.1M phosphate buffer; pH 7.4) and their midbrains post-fixed for 24h at 4°C. Thereafter, they were cryoprotected in increasing grades of buffered sucrose (10%, 20%, and 30% sucrose in 0.1M PB). Coronal midbrain cryosections of 40μm thickness were collected serially on gelatinized slides. Every 6^th^ midbrain section was chosen for immunostaining (Vidyadhara et al., 2017).

A sequential staining protocol was performed to study the proteins expression alongside co-labelling with TH (Alladi, et al., 2010a, Vidyadhara et al., 2017). Briefly, the sections were equilibrated in 0.1M PBS (pH 7.4) for 10 min and then blocked for 1h in 3% buffered bovine serum albumin (BSA). Thereafter, the sections were incubated with the primary antibody for 48 hr at 4°C. This was followed by washing and incubation with appropriate secondary antibodies i.e., Cy3-conjugated (1:200; C2821 or C2181; Sigma Aldrich, USA), FITC-conjugated (1:200; F7367 or F7512) or Cy5-conjugated secondary antibodies (1:200; AP192SA6; Merck Millipore, USA) overnight at 4 °C. Similar immunolabeling steps were followed for the subsequent sequential staining. Negative controls meant sections incubated with dilution buffer *per se*. The details of primary and secondary antibodies are provided in table 1. All the primary antibodies were sourced from Santa Cruz Biotechnology Inc, Dallas, USA and secondary antibodies from Sigma-Aldrich USA.

**Table: 1.** List of antibodies used for immunofluorescence labelling.

| Sl. No. | Primary protein | Dilution | Catalogue no. | Secondary antibody & Dilution | Source (Catalogue no.) |
| --- | --- | --- | --- | --- | --- |
| 1 | Bax | 1:200 | SC-526 (P-19) | 1:200<br>Anti-Rabbit IgG (whole molecule)-FITC produced in sheep, affinity isolated antibody adsorbed with human IgG | F7512<br>Sigma Aldrich |
| 2 | Bcl-2 | 1:200 | SC-492 (N-19) | 1:200<br>Anti-Rabbit IgG (whole mount)<br>Cy3 conjugate antibody developed in sheep | C2181<br>Sigma Aldrich |
| 3 | TH | 1:500 | SC-25269 (F-11) | 1:200<br>Donkey Anti-Mouse IgG, Alexa Fluor® 647 Conjugate | AP192SA6<br>Merck<br>Millipore,<br>United State |
| 4 | AIF | 1:200 | SC-9416 (D-20) | 1:200<br>Anti-Goat IgG (whole molecule)-FITC produced in rabbit | F7367<br>Sigma Aldrich |
| 5 | α-synuclein | 1: 500 | SC-7011-R (C-20) | 1:200<br>Anti-Rabbit IgG (whole mount)<br>Cy3 conjugate antibody developed in sheep | C2181<br>Sigma Aldrich |
| 6 | GFAP | 1:500 | SC-6170 | Anti-GFAP Antibody (C-19) | Santa Cruz<br>Biotechnology,<br>USA |

#### Image digitization by confocal microscopy

The immunofluorescently labelled images of SNpc were captured using laser scanning confocal microscope with 20X objective and magnified twice with optical zoom (DMIRE-TCS Leica, Germany). The fluorescence intensity levels were the ‘measure’ of protein expression, using the in-built software (Alladi et al., 2010b; Vidyadhara, et al., 2016a). At least 300 immunoreactive neurons per animal were sampled to derive a cumulative mean. The excitation frequency of different lasers were as follows: 488nm for FITC (AIF or Bax), 514nm for Cy3 (Bcl-2) and 633 nm for Cy5 to probe TH. Emission band widths of 495–540nm for FITC, 550–620nm for Cy3, and 649-666 nm for Cy5 were maintained to avoid non-specific overlap of emission frequencies. All the nigra images were captured using 20X objective and striatal images with 10X objective at a constant PMT voltage of 537. Other features such as optical zoom, scan speed, pinhole aperture, image resolution etc. were uniform for all observations (Alladi, et al., 2010a & b).

### Western blotting

The mice were sacrificed by cervical dislocation and the midbrains were snap frozen in liquid nitrogen. The midbrains, encompassing the nigral coordinates were cryo-sectioned, without freezing medium, into 4μm thick slices and collected in mammalian lysis buffer (Sigma-Aldrich, USA) with protease inhibitor (Alladi et al., 2002; Vidyadhara et al., 2017). The tissue was sonicated (Q Sonica, India) and protein estimated by Bradford method (TECAN infinite M200 PRO, Switzerland). 40μg of protein was loaded per lane on 12% SDS-PAGE gels, electrophoresed (Mini-protean, Bio-Rad, USA) and transferred to PVDF membranes.

The membranes were blocked using 5% skimmed milk protein in 1X PBS. The proteins of interest were Bax (1:200; SC526), Bcl-2 (1:200; SC492) and AIF (1:400; ab32516). The secondary substrate was horse radish peroxidase tagged anti-rabbit secondary antibody and anti-goat secondary antibody (1:2500, EMD Millipore). The bands were detected using a chemiluminescent substrate (SuperSignal West Pico, USA) on a gel-doc apparatus (Syngene International Ltd, India). The band intensity was estimated using Image J 1.48v program. For all the experiments, β-actin was used as the loading control.

### Complex I (CI) and Complex IV (CIV) assays

The crude mitochondrial extract was derived from each specimen (total n=24) using a mitochondria isolation kit ab110168; (Abcam UK). Briefly, the midbrain tissue was homogenized with 35 dounce strokes and spun at 1,000 g for 10 min at 4°C. The supernatant spun at 12,000 g for 15 min at 4°C was pelleted and washed twice in isolation buffer and finally re-suspended in the same buffer. After determining the protein concentration, OXPHOS activities were estimated. Activity assays were conducted using 50ug of each sample. Complex I activity was determined by following the oxidation of NADH to NAD+ and the simultaneous reduction of the dye (ε= 25.9/mM/well); measured at OD 450 nm by using Complex I Enzyme Activity Microplate Assay Kit (Colorimetric) (ab109721; Abcam UK). Complex IV which was immunocaptured within the wells was assayed for the activity by colorimetric determination following the oxidation of reduced cytochrome c as decrease in absorbance at 550 nm.

### Electron Microscopy

The mice were anesthetized (n=3-4 per strain/experimental condition; total 32) with halothane inhalation and perfused transcardially with a mix of glutaraldehyde (2.5%) and paraformaldehyde (PFA 2%) in 0.1M phosphate buffer (PB, pH 7.2) for 3 min. After fixation, they were processed in 1% osmium tetroxide for 1 hr at 4°C. The tissues were dehydrated through an ethanol series (70%, 80%, 90%, 95%) for about 1 hr in each solution followed by two changes of absolute alcohol for 30 min. The specimens were cleared twice in propylene oxide to facilitate infiltration and embedded in resin using araldite CY212 mixture. The tissue samples were then embedded in resin using beam capsules and allowed to polymerize at 60° C for two days. After polymerization, 1μm semi thin sections were collected on slides and stained with 1% toluidine blue to observe the region of interest and orientation of cells. Ultrathin (80 nm) sections were cut using UC6 ultramicrotome (Leica, Austria) and mounted over copper grids. The sections were stained with saturated uranyl acetate for 1 hr followed by 0.2% lead citrate (5-7 min) for better image contrast. The sections were washed and then air dried for few minutes and examined under transmission electron microscope (TECNAI G2 Spirit BioTWIN, FEI, Netherlands).

### Fecal microbiome analysis

Bacterial 16S rRNA hypervariable regions V3-V4 were extracted and amplified from the fecal matter of all mice strains (n=3/strain; total n=12) using V3V4F and V3V4R primers (Clevergene Biocorp Pvt. Ltd, Bangalore). Briefly, DNA was extracted from mice fecal pellets using PureLink Microbiome DNA Purification kit (Invitrogen, USA) following standard protocol. Bacterial 16S rRNA hypervariable regions V3-V4 were amplified using V3V4F and V3V4R primers (Klindworth et al., 2013). 25ng of DNA was used for PCR amplification using KAPA HiFi HotStart Ready Mix. The amplicons were purified using Ampure beads to remove unused primers. Libraries were quantitated using Qubit dsDNA High Sensitivity assay kit. Sequencing was performed using Illumina Miseq with 2×300PE V3 sequencing kit. The sequence data quality was checked using FastQC and MultiQC software. The data was checked for base call quality distribution, % bases above Q20, %GC, and sequencing adapter contamination.

The QC passed reads were imported (Schloss et al., 2009) and the pairs were aligned with each other to form contigs. The contigs were screened for errors and only those between 300bp and 530bp were retained. Finally, the filtered contigs were classified into taxonomical outlines based on the Silva v.128 database (Quast et al., 2013) and were clustered into OTUs (Operational Taxonomic Unit). After the classification, OTU abundance was estimated. OTUs with abundance less than 0.01% of the total reads were identified as singletons or rare OTUs and were removed from all downstream analysis. A total of 196 distinct OTUs with at least 100 reads supporting each were identified.

### Statistical Analysis

The data was analysed using 2-way ANOVA followed by Tukey’s post hoc test. The values were expressed as mean + SD and a p-value lower than 0.05 was considered significant.

## Results

### Apoptosis Inducing Factor (AIF) is up-regulated following MPTP administration

AIF expression was pan cytoplasmic in the dopaminergic neurons. Amongst the adults, the pure strains had higher baseline AIF expression as compared to the crossbreds. The F1X2 had significantly low AIF expression (Figure 1. A & B *p <0.05 CD-1 vs. F1X2). Following MPTP administration, its expression in the C57BL/6J increased significantly compared to the rest (Figure 1.B ****p<0.0001 C57BL/6J-Normal vs. MPTP, *p<0.05 F1X1-Normal vs. MPTP; ***p<0.001 F1X2-Normal vs. MPTP).

**Figure 1.**
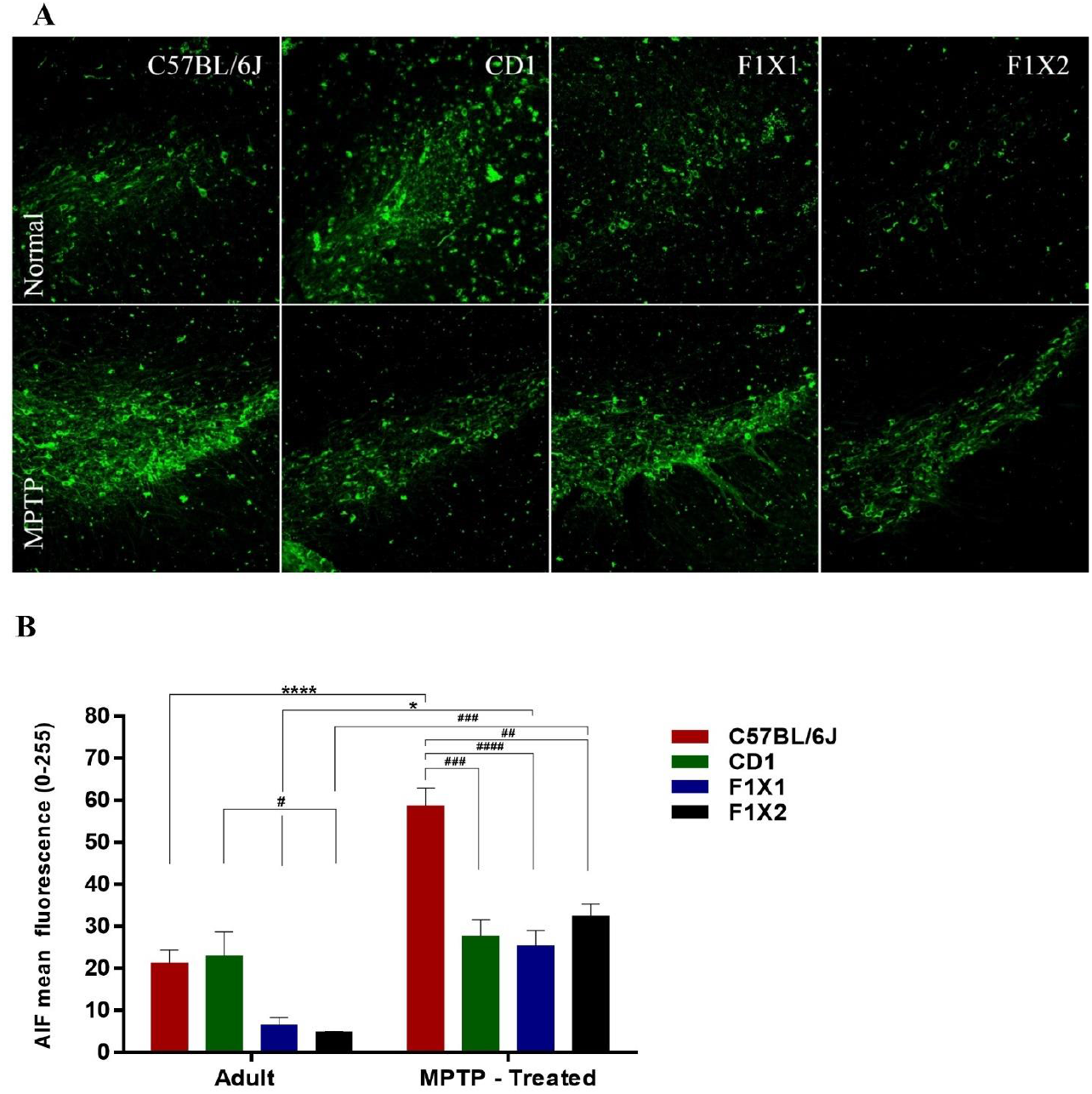
Representative immunofluorescent images showing staining of AIF in DA neurons of the SNpc (A). There was a significant upregulation of AIF expression following MPTP administration (B) ****p<0.0001 C57BL/6J. Normal vs. MPTP, *p<0.05 F1X1-Normal vs. MPTP, ***p<0.001F1X2-Normal vs. MPTP. Western blots also showed similar trends of AIF expression as seen in the densitometry quantification but were not statistically significant (Figure 6).

### MPTP alters expression pro- and anti-apoptotic proteins of BCL-2 family

Both the DA and non-DA midbrain neurons expressed the pro-apoptotic (Bax) as well as anti-apoptotic protein (Bcl-2). The TH immunoreactive DA neurons in the SNpc that expressed Bax and, Bcl-2 were considered for quantification. Both the proteins were prominently perinuclear in localisation (Figure 2A). The baseline Bax and Bcl-2 expression was low in the crossbreds as compared to C57BL/6J or CD-1 (Figure 2B. Bax. **p < 0.01 C57BL/6J vs. F1X1, ***P < 0.001 F1X2, *p < 0.05 CD-1 vs. F1X1; C. Bcl-2. ***P < 0.001 C57BL/6J vs. F1X2). In response to MPTP, Bax expression was significantly up-regulated, whereas Bcl-2 remained unaltered. The Bax: Bcl-2 ratio, increased in response to MPTP in all the groups (Figure 2.D. *p < 0.05 normal vs. MPTP-C57BL/6J, *p < 0.05 normal vs. MPTP-CD-1, **p < 0.01 normal vs. MPTP-F1X1, **p < 0.01 normal vs MPTP-F1X2).

**Figure 2.**
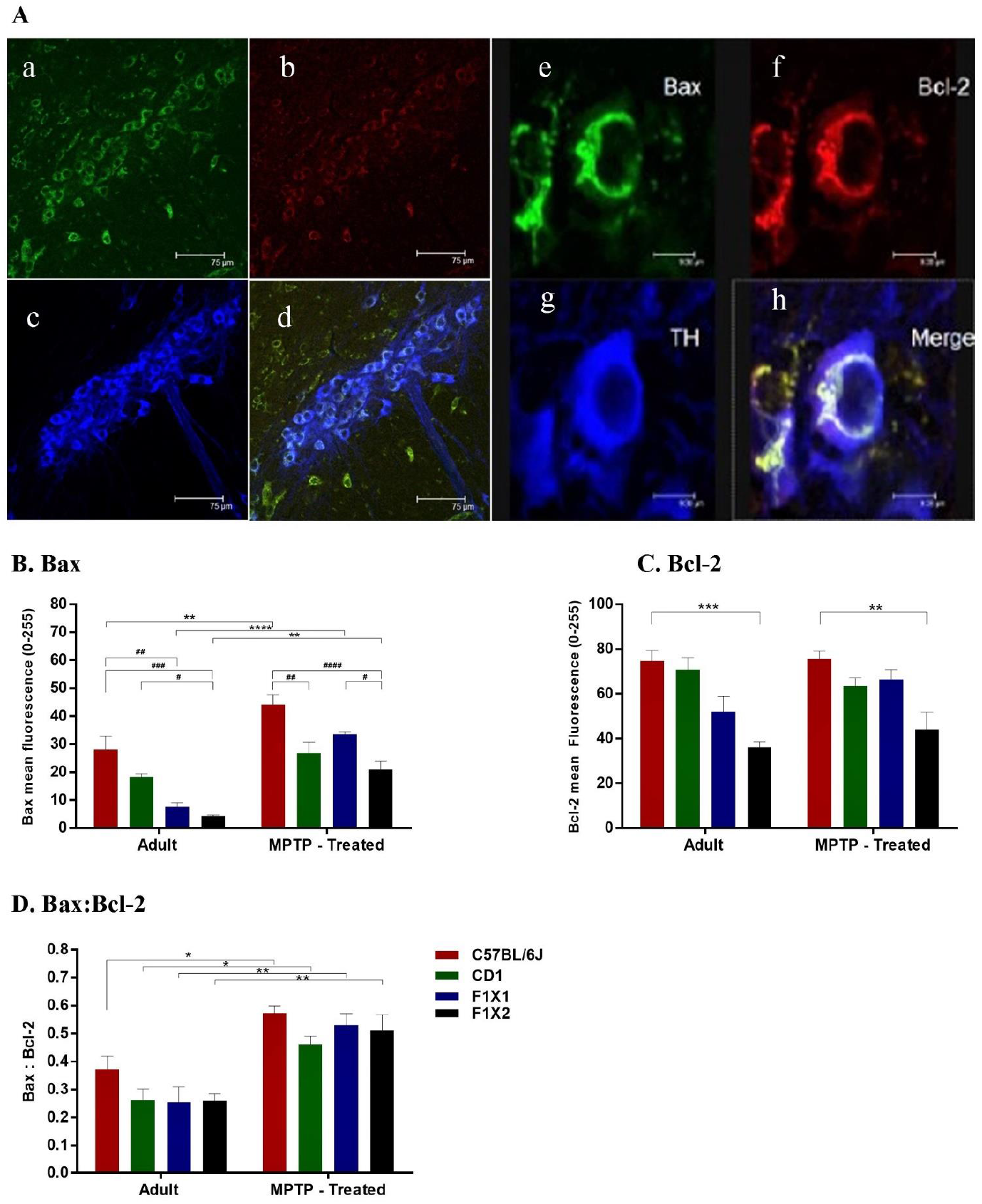
Laser scanning confocal micrograph showing triple-labeling immunostaining of Bax (A.a), Bcl-2 (A.b), and TH (A.c) in dopaminergic neurons (A.d. merged). The proteins Bax (A.e) and Bcl-2 (A.f) are localized to the cytoplasm and are more intense in the perinuclear region (A.h). Bax and Bcl-2 expression in the normal groups was relatively low in the crossbreds when compared to C57BL/6J or CD-1 (Figure 1.B. Bax **p < 0.01 C57BL/6J vs. F1X1, ***P < 0.001 C57BL/6J vs. F1X2, *p < 0.05 CD-1 vs. F1X1; C. Bcl-2 ***P < 0.001 C57BL/6J vs. F1X2). In the MPTP injected mice, Bax was significantly up-regulated, while, there was no change in the levels of Bcl-2. Note the relatively high Bax:Bcl-2 ratio in the C57BL/6J and increased following MPTP administration in all the groups (Figure 5.D. Bax:Bcl-2 *p < 0.05 C57BL/6J Normal vs. MPTP, *p < 0.05 CD1 Normal vs. MPTP, **p < 0.01 F1X1 Normal vs. MPTP, **p < 0.01 F1X2 Normal vs MPTP).

### MPTP induced sizeable degeneration in C57BL/6J

A semiquantitative evaluation of AIF, Bcl-2 and Bax in the midbrain tissue immunoblots, showed significant up-regulation of AIF and pro-apoptotic protein Bax following MPTP exclusively in the C57BL/6J mice (Figure 3B. *p < 0.05 C57BL/6J Normal vs MPTP; D. **p < 0.01 C57BL/6J Normal vs MPTP). The other strains showed mild, non-significant alterations (Figure 3C & E).

**Figure 3.**
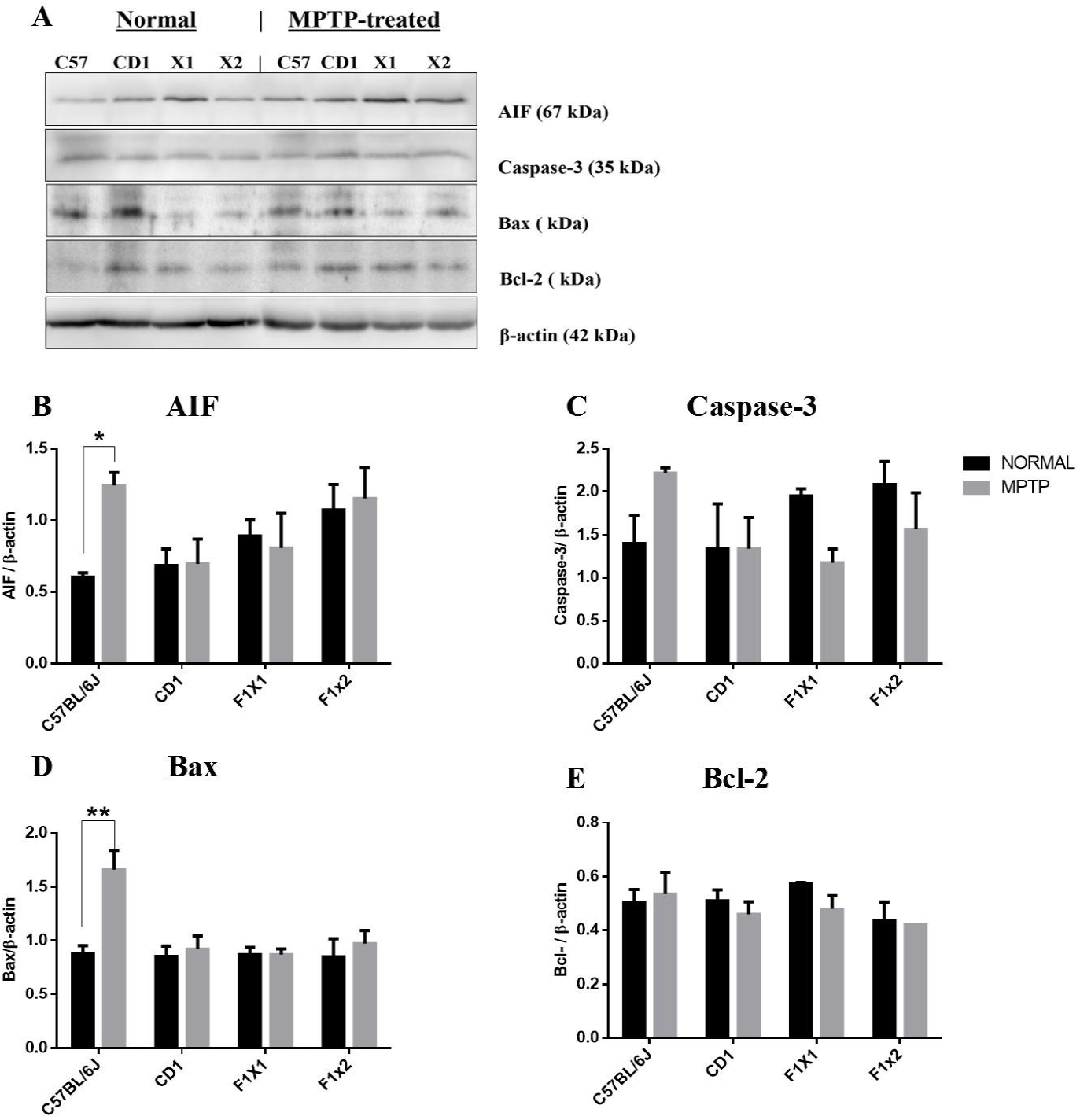
Apoptotic protein expressions increased upon exposure to MPTP. Representative immunoblots showing basal AIF, Caspase-3, Bax and Bcl-2 expression in normal and MPTP-administered mice. MPTP treatment significantly increased AIF and Bax levels in C57BL/6J (B. *p<0.05 C57BL/6J normal v/s MPTP-injected; D. **p<0.01 C57BL/6J normal v/s MPTP-injected) whereas it was maintained in CD-1 and crossbreds. Note the slight increase in Caspase-3 in C67BL/6J while it decreased in the F1X1 and F1X2 upon MPTP administration (C). Bcl-2 expression did not show any significant differences between the two groups (D).

### MPTP causes an increase in α-synuclein expression in the pars reticulata of F1X2

Amongst the parent strains, C57BL/6J showed higher α-synuclein expression than CD1. The basal α-synuclein levels in SNpc were comparable across the four mice strains studied. Following MPTP injection too, no significant changes were noted in the expression levels across the different strains (Figure 4A). Surprisingly, an increase in α-synuclein expression was noted in the SNpr of both the crossbreds, post-MPTP treatment (*p<0.05; Fig 3).

### Expression of striatal GFAP is associated with GDNF

In our earlier study, we noted an increase in the striatal GDNF expression in the resistant strains (Vidyadhara et al., 2017). Since GDNF is referred as “astroglia derived”, we assessed GFAP expression in the striatum. GFAP expression was similar in all the strains under normal conditions. Following MPTP administration, GFAP was significantly upregulated, exclusively in the crossbreds (Figure 5. A & B. *p < 0.05 normal vs. MPTP-F1X1, ***p < 0.001 normal vs. MPTP-F1X2). In F1X2, the increase in GDNF expression (Figure 5B and L) could be linked to GFAP expression (Figure 5L), which lacked a little in the F1X1.

**Figure 4.**
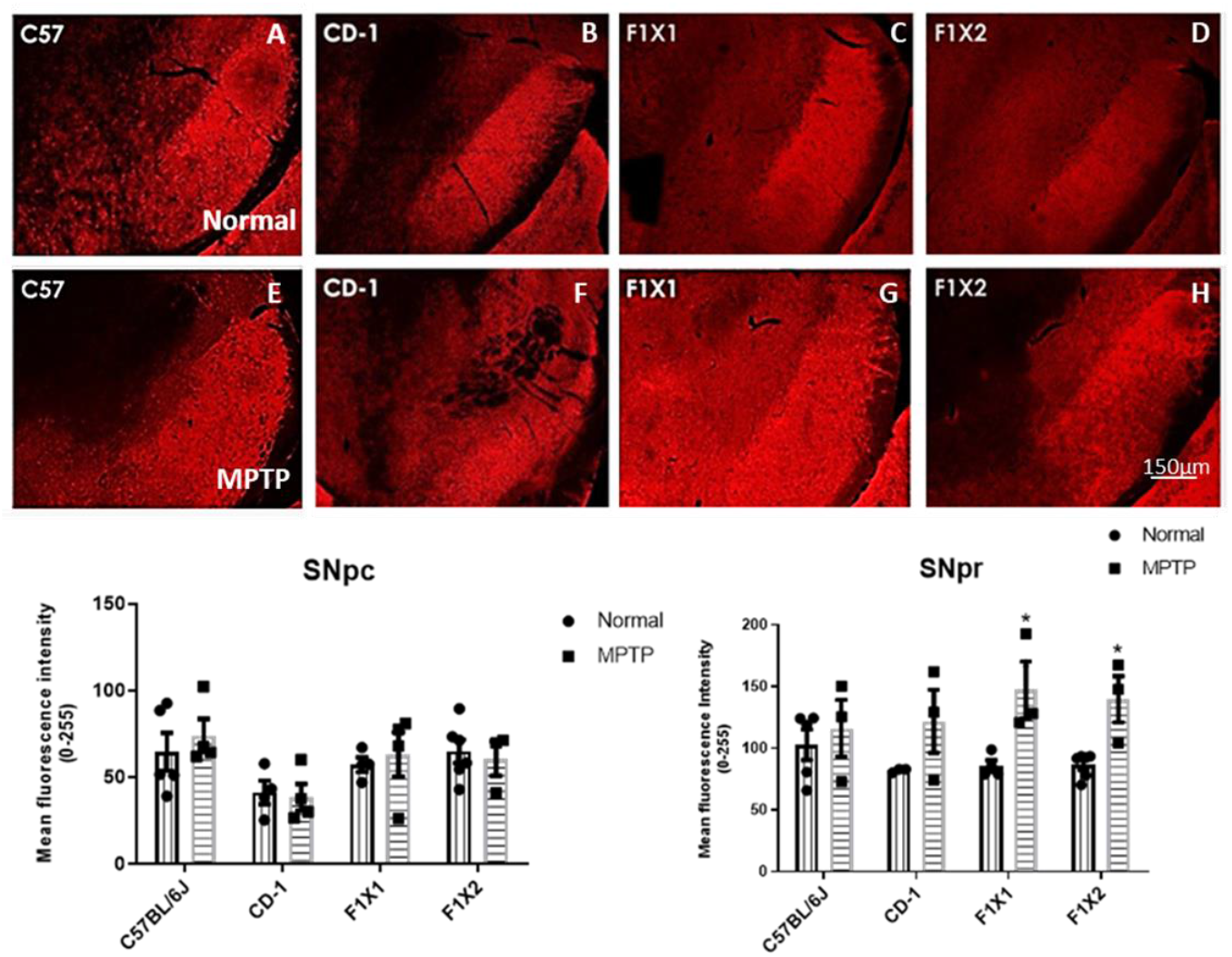
Representative immunofluorescent images showing staining of α-synuclein in the SNpc and SNpr. Note the expression is associated with the neuropil of the pars reticulata is increased in the F1 crossbreds following MPTP.

**Figure 5.**
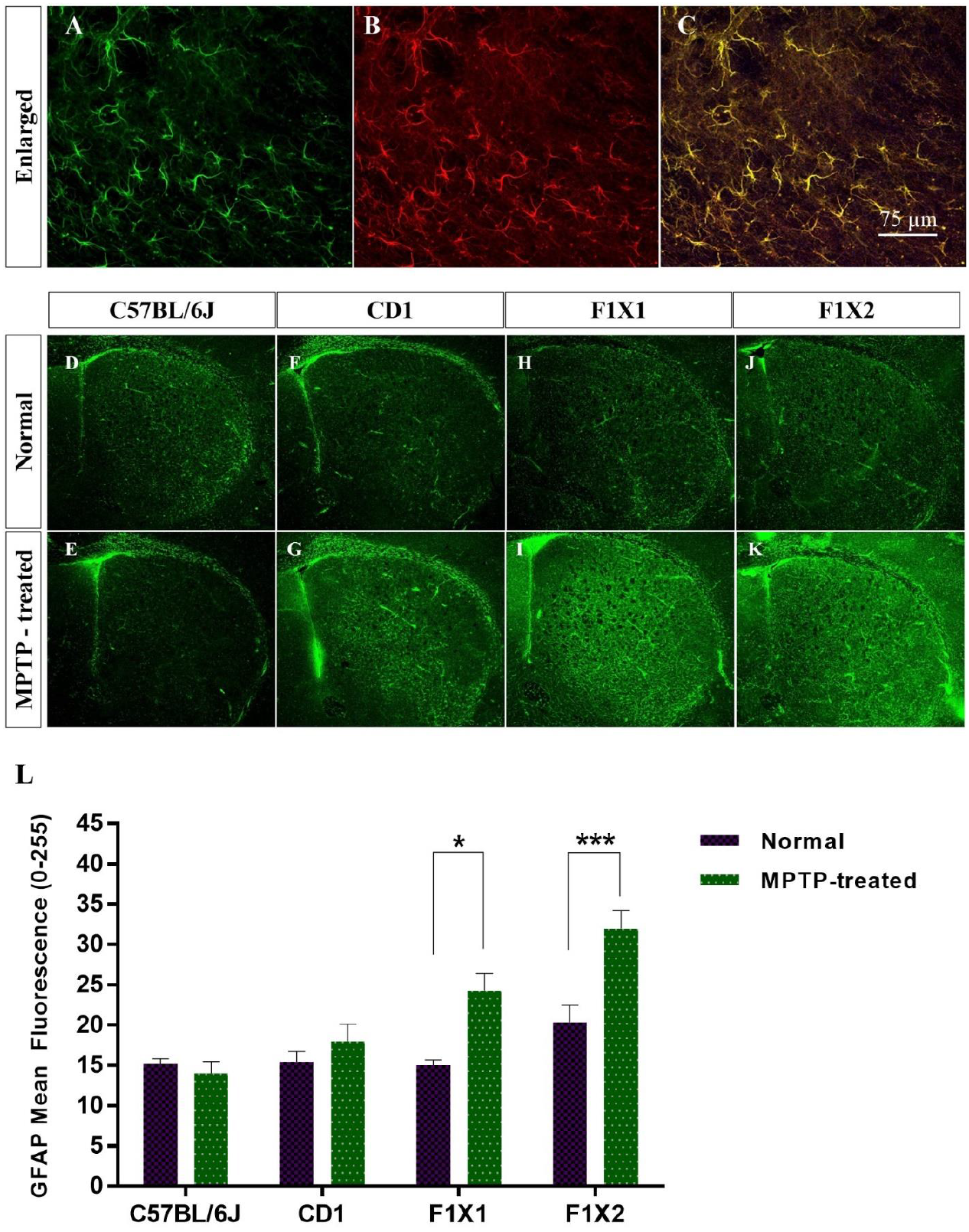
Immunofluorescence staining for GFAP in the striatum in normal and MPTP injected groups. The higher magnification image reveals co-expression of both GFAP (A. Green) and GDNF (B. Red). In the experimental group, only F1X1 and F1X2 (H, I, J, K and L) showed a significant upregulation of GFAP expression in the striatum while there was no difference in the control groups (D, E, F G and L *p < 0.05 normal vs. MPTP-F1X1, ***p < 0.001 normal vs. MPTP-F1X2). Scale bar 75 μm.

### Mitochondrial complex-I activity was decreased in the susceptible strain

Mitochondrial complex-I activity decreased in C57BL/6J following MPTP administration (Figure 6A. *p< 0.05 C57BL/6J Normal vs. MPTP), whereas in CD-1 and the crossbred mice, it remained unaltered. Interestingly, we saw an increase in the complex-IV activity in C57BL/6J and CD-1 (Figure 6B. **p< 0.01 C57BL/6J-Normal vs. MPTP; **p< 0.01 CD-1-Normal vs. MPTP). The crossbreds, particularly, the F1X2 showed no alterations in their mitochondrial complex activities in response to MPTP.

**Figure 6.**
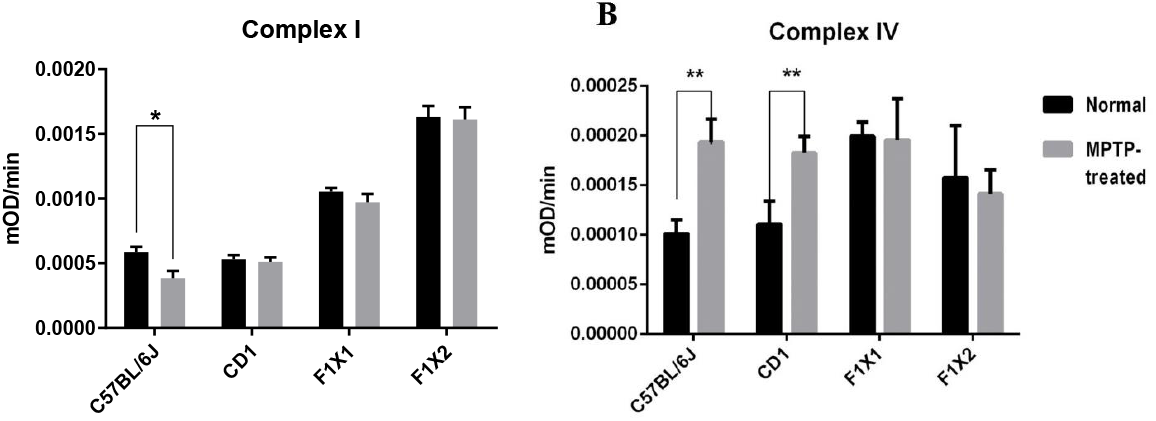
Histograms showing mitochondrial complex-I (A) and complex-IV (B) activity in control and MPTP-challenged mice midbrains. Note the decreased complex-I activity following MPTP-injection in C57BL/6J (Figure 13.A. **p< 0.01 C57BL/6J Normal vs. MPTP). We saw a compensatory increase in complex-IV function (Figure 13.B. **p< 0.01 C57BL/6J Normal vs. MPTP; **p< 0.01 CD-1-Normal vs. MPTP). 50 μg of crude mitochondrial extract were utilized to test the activity using Abcam complex assay kits (ab109721and ab109911)

### MPTP induced ultrastructural changes

The ultrastructural features of the DA neurons in the control groups were well appreciable, in form of intact Golgi apparatus, both round and elongated mitochondria and long continuous ER. The axo-synaptic synapses and vesicles were quite common too (Figure 7A-D, Normal). MPTP caused severe disintegration of DA neurons in C57BL/6J, on the contrary the CD-1 nigra harboured partly affected and healthy neurons. In C57BL/6J, the mitochondrial condensation as well as Golgi complex disruption was noted. Qualitative observations showed a decrease in the number of ribosomes on the rough ER, and an increase in the number of free polyribosomes (Figure 7, compare A vs. E). The CD-1 neurons showed more long tubular mitochondria, preserved Golgi apparatus, but dilated ER. The F1X1 and F1X2 had remarkably preserved organelles. There appears to be no synaptic disturbances following MPTP injection, as evinced by a steady density of synaptic vesicles, comparable in control (Figure 7 A-D) and MPTP-injected mice nigra (Figure 7 E-H).

**Figure 7:**
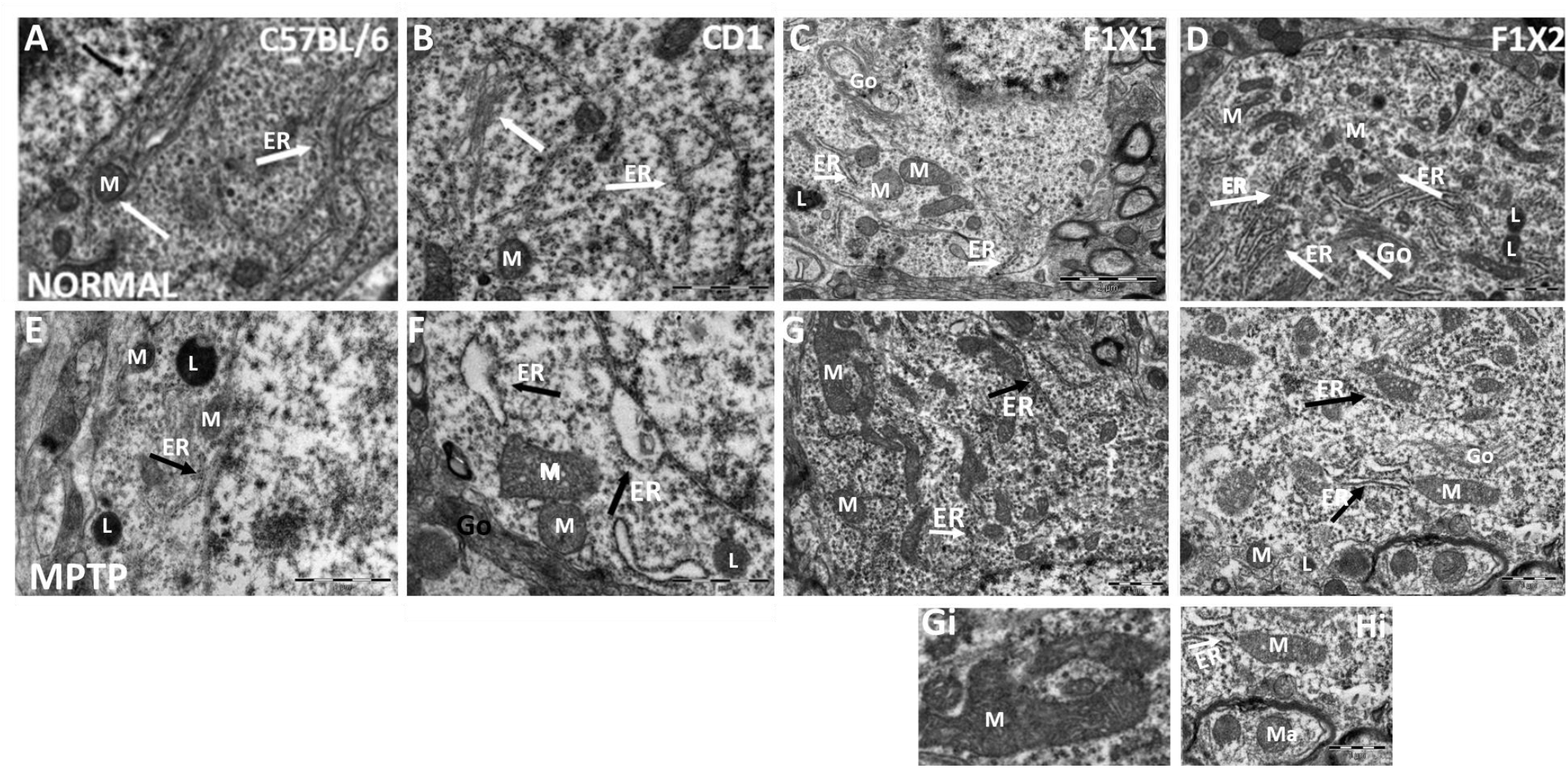
Representative transmission electron micrographs showing ultrastructural features of the DA neurons in the normal mice (A-D) were well appreciable, in form of intact Golgi apparatus, both round and elongated mitochondria (M) and long continuous ER (White arrows, A-D, Normal). Note the MPTP induced ultrastructural changes (E-H). Note the presence of small mitochondria (M) and fragmented ER (ER & black arrows) in MPTP injected C57BL/6J. On the contrary the CD-1 nigra harboured healthy and partly affected (F) neurons. Note the dilated ER, normal sized mitochondria (M). Qualitative observations showed a decrease in the number of ribosomes on the rough ER, and an increase in the number of free polyribosomes (Compare A&B with E&F). (C&G) The normal nigral DA neurons of F1X1 mice show normal mitochondria, Golgi and elongated ER. (G) Note the presence of normal and few small ER, and increased ribosomes. Following MPTP, the mitochondria included small and elongated tubular ones (Gi). (D&H) The DA neurons of normal F1X2 show intact Golgi apparatus (Go), round and elongated mitochondria, elongated ER (Control, D). In the MPTP injected group, normal and elongated (M) as well as axonal mitochondria (in Hi: Ma) were noted. Note both normal and dilated ER (MPTP, H; Hi).

Although MPTP failed to induce apoptosis in all the DA neurons of SNpc, it certainly had detrimental effect on the organelles. The maximum ultrastructural injury was observed in the susceptible C57BL/6J; though the neurons in the resistant strain were also affected. Since there was no large-scale loss of nigral neurons in response to MPTP in the crossbreds, the moderate level of ultrastructural alterations may reflect sub-threshold degeneration in the F1 crossbred specimens.

### Synapses within the neuropil of SNpc in different mice strains

Representative electron micrographs of synapses in the control (A) C57BL/6J and (B) CD-1 (C) F1X1 appear to have several synaptic vesicles (Sv) at the presynaptic site and electron dense post synaptic densities (box). Note the presence of series of synapses in the F1X2 D, box; arrow), with vesicles and several mitochondria in the presynaptic site (***). There were no perceptible alterations in the MPTP-injected nigra of these mice (D-G). The MPTP-injected crossbreds too harbor synapses that are slightly less electron dense, but preserved (Compare C vs. G and D vs. H). Insets show high mag images of the synapses of the respective study group.

### The resistant strain and crossbreds harbor more identical gut microbiome

Of the many genera that showed on the heat map distribution the top 10 of the bacterial genus were picked and we correlated their abundance in different groups. The gut microbes in C57BL/J had more genera belonging to the Streptococcus, Bacteroides and Lachnospiraceae family whereas its presence is relatively lower in CD-1 and the crossbreds (Figure 9). The neurotoxin resistant strain CD-1 and the crossbreds seemed to harbor more microbes and particularly in abundance were those belonging to Prevotellacea family. We observed the maximum concentration of bacteria of the Prevotella spp in F1X2.

**Figure 8:**
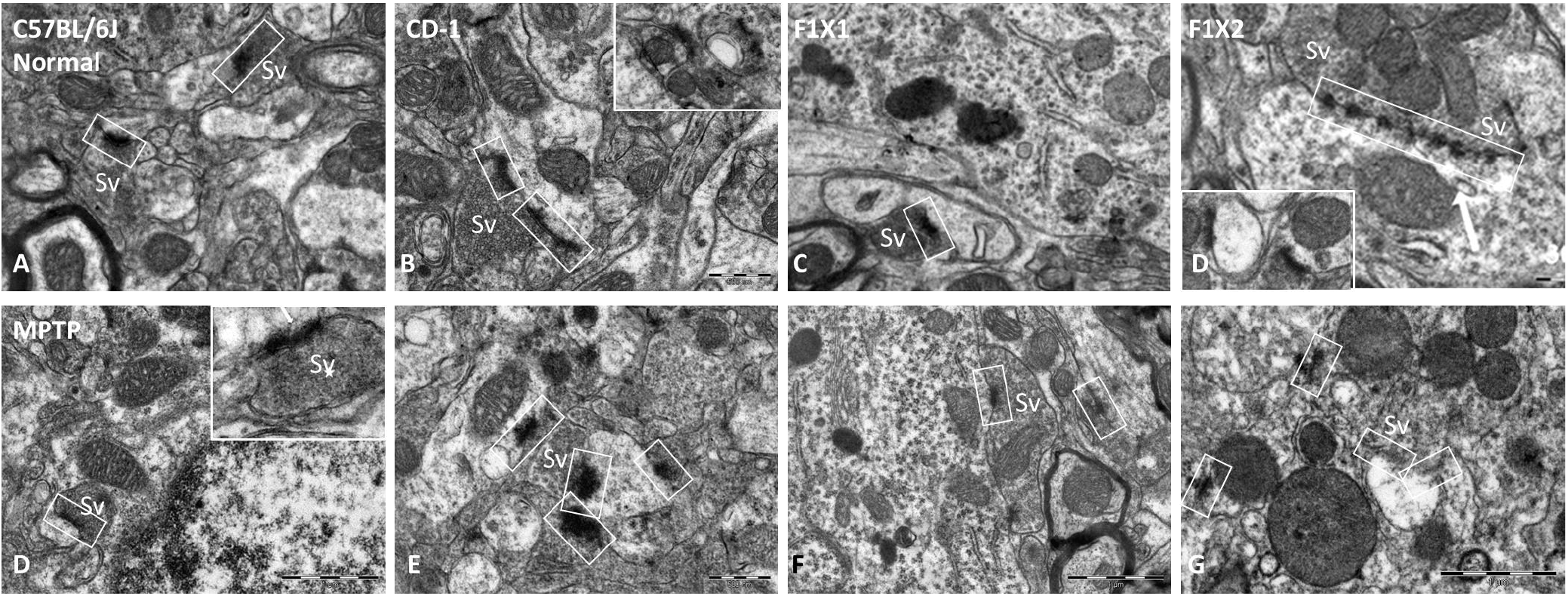
Synapses within the neuropil of SNpc in different mice strains. Representative electron micrographs of synapses in the control (A) C57BL/6J and (B) CD-1 (C) F1X1 appear to have several synaptic vesicles (Sv) at the presynaptic site and electron dense post synaptic densities (box). Note the presence of series of synapses in the F1X2 D, box; arrow), with vesicles and several mitochondria in the presynaptic site (***). There were no perceptible alterations in the MPTP-injected nigra of these mice (D-G). The MPTP-injected crossbreds too harbor synapses that are slightly less electron dense, but preserved (Compare C vs. G and D vs. H). Insets show high mag images of the synapses of the respective study group.

**Figure 9.**
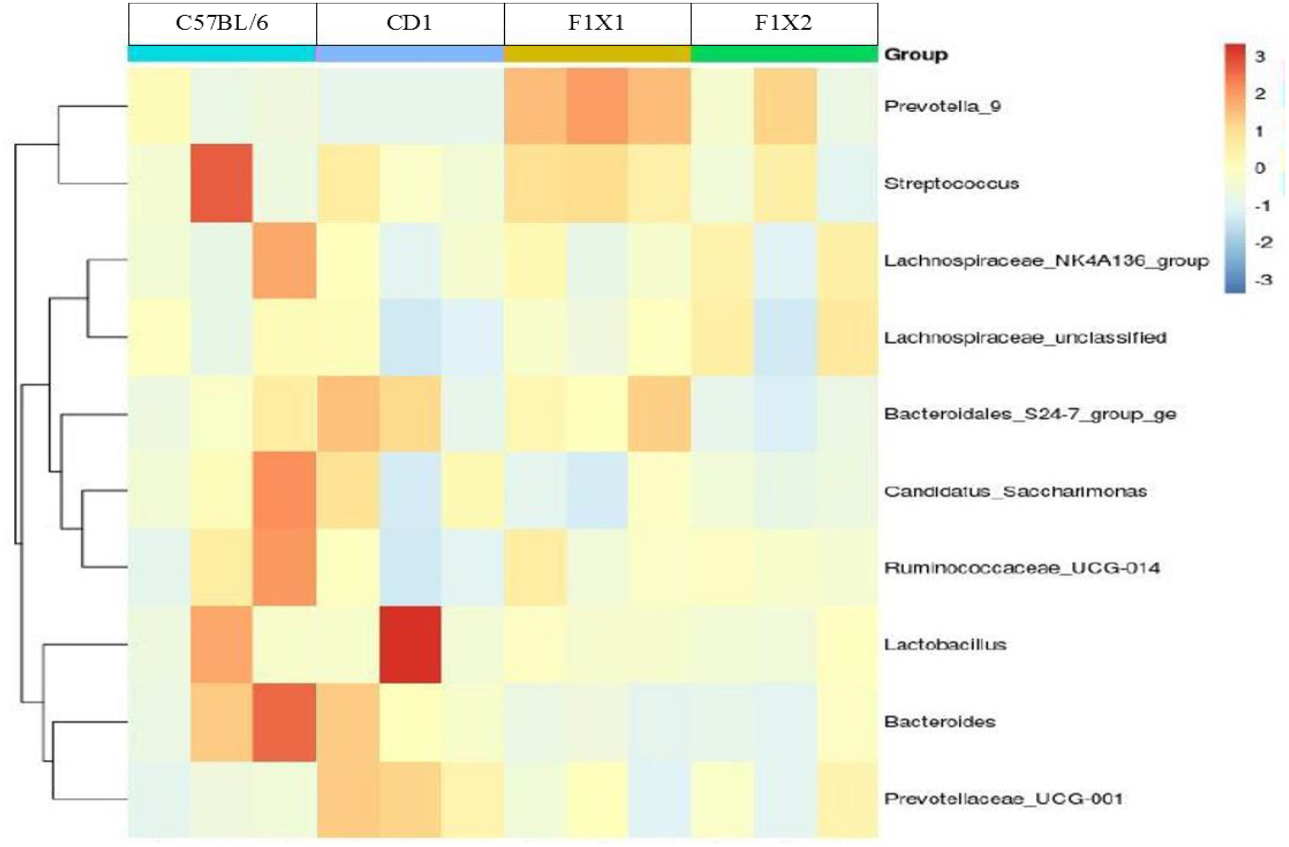
Heat map representing top 10 Genus of gut microbiota following metagenomic analysis from the fecal matter. The F1X1 and F1X2 crossbreds showed the highest abundance of bacteria as compared to C57BL/6J or CD-1 including those Prevotellacea and Lachnospiraceae which are reportedly shown to be reduced in human PD patients when compared to normal subjects.

## Discussion

### Enhanced AIF, may underlie enhanced susceptibility and cell death

We evaluated the molecular mechanisms/factor other than caspase-mediated cell death in the MPTP-induced neurodegeneration. AIF is a mitochondrial flavoprotein located in the intermembrane space and upon its release in response to stimuli. It participates in the intrinsic apoptosis pathway (Eliasson et al., 1997) by causing chromatin condensation and DNA fragmentation. Upon sensing death signals, AIF is released from the mitochondrial outer membrane and is translocated to nucleus where it regulates Parkin assisted neuroprotection (Guida et al., 2019) or cell death as seen in traumatic brain injury, oxidative, cerebral hypoxia–ischemia stress, epilepsy and Alzheimer’s disease (Chu et al., 2005; Hegde et al., 2002). The increased AIF expression in the MPTP-administered groups in all the strains emphasizes the role of caspase-independent cell death mechanisms, in degeneration of dopaminergic neurons in mouse nigra. AIF along with caspase-3 co-operatively mediate apoptosis in T-cell lineage (Prabhu et al., 2013). Interestingly, the C57BL/6J showed maximum upregulation re-emphasizing its vulnerability, compared to the resistant CD-1 or the crossbreds. The congruent action of AIF and caspase may yield the common apoptotic phenotypes such as neuronal shrinkage noted in our earlier study (Vidyadhara et al., 2017). AIF may possibly be the ‘long hypothesized, hitherto unknown non-caspase component’ that mediated the cellular commitment death. Thus, both AIF-dependent and caspase-dependent mechanisms may contribute independently and synergistically in the final execution of dopaminergic neuronal apoptosis.

### Higher Bax: Bcl-2 ratio exaggerates MPTP-induced cell death in C57BL/6J mice

Adult C57BL/6J have relatively fewer nigral neurons and show enhanced apoptosis in response to MPTP than CD-1 and their crossbreds (Vidyadhara et al., 2017). Bcl-2 is associated with MPTP-induced apoptosis pathway in DA neurons (Liu et al., 2018, Song et al., 2018; Pajarillo et al., 2018). A balance between Bcl-2 and Bax is vital to form heterodimers and to modulate the programmed cell death pathway (Certo et al., 2006; Strasser et al., 2011). Activation of JNK/SAPK and the increased ratio of Bax to Bcl-2 implores Bax to have has larger active contribution in neurodegeneration. The preserved ratio of Bcl-2: Bax in the crossbreds suggest better neurotrophic action in their nigra, ably balanced by striatal GDNF (Bekker et al., 2021) and well reflected by preserved striatal field potentials reported earlier (Vidyadhara et al., 2017; Vidyadhara et al., 2019). AIF expression as well as Bcl-2/Bax in dopaminergic neurons, suggests caspase dependent and independent mechanisms being at fray in the mice nigra. Accordingly, the C56BL/6J appears to be more vulnerable to minimal toxic insults, when compared to the other strains.

### Mild astrogliosis supports DA neuronal survival upon MPTP Administration

Some molecular factors associated with DA neuronal survival and apoptosis identified include GDNF and the Bcl-2 family. GDNF is a promising therapeutic candidate for PD, as demonstrated in a mice model (Espinoza et al., 2021). The co-localisation of GDNF with GFAP in the striatum in our study, validates the astrocytic origin of the trophic factor. The concomitant increase in both proteins in the F1X2 crossbreds attribute a positive role for astroglia. Upregulation of astroglia is considered to be safer amongst the modalities of GDNF deliveries being currently explored (Patel et al., 2019).

#### Over expressed α-synuclein in reticulata, an indicator of sub-threshold neurodegeneration

α-synuclein is a key regulator of synaptic vesicle release and trafficking, neurotransmitter release, dopamine metabolism and release, fatty acid-binding, and neuronal survival. It is central to the pathogenesis of PD. Over-expression of the physiological α-synuclein, as detected via the immunofluorescence assay, is considered detrimental to the neuronal function. In a similar parlance, in a US-based study, the presence of Lewy pathology and logarithmic increase in physiological α-synuclein in aging human nigra, was linked to a pre-Parkinsonian state (Chu et al., 2007). Contrarily, we found a gradual non-logarithmic increase in α-synuclein as well as the presence of the Lewy pathology only after 6^th^ decade in the aging substantia nigra of the Asian-Indian cohort. The latter was much subdued compared to the US cohort, and was attributed to the lower prevalence to PD in the Asian-Indian populations (Alladi et al., 2010b).

Indeed, Lewy body pathology, comprising of oligomeric and fibrillar pathogenic α-synuclein, is not formed in the acute MPTP mice models, unlike, the MPTPp (MPTP + Probenecid) which shows more gradual and progressive DA neuron loss, resulting from pathogenic accumulation and aggregation of α-synuclein. Thus, the lack of alterations in α-synuclein expression in the SNpc of different mice strains as well as in response to MPTP may be due to the acute MPTP regime followed in our study.

The par reticulata comprises of GABAergic neurons as well as the anatomically deflected dendrites of the pars compacta moving ventrally. The dendrites release dopamine, which by the virtue of being extrastriatal, may modulate activity-dependent synaptic plasticity of the output neurons of the reticulata. The enhanced α-synuclein expression in response to MPTP in the pars reticulata of crossbreds may hint at enhanced neurotransmission and hence a neuroprotective stratagem. Alternatively, it may herald sub-threshold degeneration of the dendrites of DA neurons or the SNpr *per-se*. α-Synuclein being a synaptic protein, aberrant synaptic plasticity in SNpr, may contribute to pathophysiology of PD (Prescott et al., 2009). Faynveitz et al., (2019) demonstrated that 6-OHDA-induced depletion of dopamine in the SNpc neurons, results in reduced firing rate and periodicity of the SNr neurons, suggesting expansion of GABAergic inhibition within the SNr, and thus contributing to PD pathology.

### Ultrastructural damage and mitochondrial complexes activity in MPTP-induced neurodegeneration

Aberrations in mitochondrial morphology were described in the tissues of PD patients (Tanaka et al., 1988). Experimental tissues of dog, and monkey (Rapisardi et al., 1990) describe analogous mitochondrial anomalies viz. quantitative reduction, membrane disruption, intra-mitochondrial paracrystalline inclusions and mitochondrial swelling as well as abnormal distribution involving formation of small clusters (Anglade et al., 1997; Forno & Norville, 1976; Trimmer et al., 2000). In our observations, the qualitative decrease in number of ribosomes on the RER were complemented by increase in the number of free polyribosomes probably suggesting an insult-induced compensatory priming of protein synthesis machinery. In the SNpc of MPTP exposed C57BL/6J mice, we found similar abnormalities in the mitochondria (Abhilash et al., 2020; preprint), which were complemented by the expression of fission related proteins (Seshadri and Alladi 2019). The present findings parallel the previous ones, thus harnessing the reproducibility of the observations. The neuroprotective features in the resilient strains reminisce the absence of mitochondrial disturbances in curcumin pre-treated mice (Pan et al., 2012). The ultrastructural features of the DA neurons of the injected crossbreds matched that of controls e.g., with intact Golgi, both round and elongated mitochondria and long continuous endoplasmic reticulum, suggesting fairly well preserved organellar architecture. Although MPTP may not induce apoptosis in all the SNpc-DA neurons, it certainly induces few degenerative changes, reflecting as alterations in the organellar structure. In the crossbreds, in the absence of large-scale neuronal loss in response to MPTP (Vidyadhara et al., 2017), the moderate ultrastructural alterations advocate the occurrence of sub-threshold neurodegeneration.

The SNpc receives glutamatergic inputs from the subthalamic nucleus and the pendunculo-pontine nucleus (Mena-Segovia et al., 2004) and GABAergic inputs from the striatum. MPTP enhances the density of Vglut2+ terminals synapsing on TH (-) dendrites within the SNpc; while the TH (+) dendrites show a significant decrease within the SNpc (Moore et al., 2021). The synapses of the MPTP injected animals in all the strains following MPTP administration were preserved, suggesting the health of the surviving neurons. It is likely that these may be the neurons that survive the MPTP-induced neurotoxicity and hence show robust synapses, or vice versa. It is also likely that the various innervations that the SNpc receives, viz. the pendunculo-pontine nucleus, STN etc. are preserved, because of the lower concentrations of MPTP used in our model compared to others (Moore et al., 2021). It might be worthwhile to explore an earlier timepoint to examine the effects on synapses.

The MPTP-induced aberrations of neuronal organelles were echoed by the dysfunction of mitochondrial complex-I activity in the susceptible C57BL/6J following MPTP administration, similar to that found in the experimental model of PD, resulting in midbrain dopaminergic neuronal loss (Meredith and Rademacher, 2011; Valente et al., 2004). In PD patient tissues, mitochondrial complex-I activity decreased in the nigrostriatal region and cortex (Schapira et al., 1989; Parker et al., 1989). The surprising finding of compensatory increase in the complex-IV activity in C57BL/6J and CD-1, suggests an effort to compensate. The stability of mitochondrial complexes in the crossbreds e.g., F1X2 is complemented by the preserved mitochondrial architecture following MPTP-injection. Thus, the DA neurons of the susceptible strain accede readily to toxic insults and are specifically defenceless against neurodegeneration.

### Gut microbiome is associated with risk for PD

Gastro-intestinal disturbances precede the motor symptoms, and environmental factors play an important role in PD. Microbial colonization occurs at the time of birth, and the cells undergo diversification and changes at infancy, to become a steady structure at about three years to retain the architecture though there are variations between individuals (Koenig et al., 2011). In PD, the composition of this complex systems has been reportedly altered (Scheperjans et al., 2015; Fung et al., 2017). The gut microbes influence the CNS through the cross-talk i.e., the “gut-brain axis”. The microbiota support the host via the secretion of SCFA (Unger et al., 2018). Impairment of the SCFA receptor FFAR2, mimic microglia injury in germ free conditions suggesting that host bacteria govern microglia, inflammation and microglial cell activation in the brain (Erny et al., 2015). The abundance of Prevotella spp and lesser Enterobacteriaceae in CD-1 and F1X2 as well as the negative association of Prevotella with disease duration in PD patients (Scheperjans et al., 2015) complement two systems. The opposite was true for the susceptible C57BL/6J which had abundance of Bacteroides and Streptococcus. This underlines the existence of an efficient interlocution between the gut bacterial community and an individual’s susceptibility/risk of suffering from PD. Although our observations on the fecal microbiome are preliminary, it is likely that few resident microbes; their secretions and metabolites may have modulated host susceptibility to diseases. Identification of such microbes and the possible bridge that links the gut and the brain with respect to PD might enable early identification of individuals at risk and provide biomarkers for disease severity and progression.

Thus, our study provides detailed mechanisms pertaining to factors associated with the differential susceptibility of mice strains C57BL/6J, CD-1 and their crossbreds to neurotoxin MPTP. The normal genetic composition of an individual or a population, sans major mutations; appears to be critical in determining susceptibility to the disease.

## Acknowledgements

The authors are grateful to Dr. G.H. Mohan, Head Veterinarian at National Centre for Biological Sciences, Bengaluru for providing breeding colonies of CD-1 mice strain.

## Ethical approval

All applicable international, national, and/or institutional guidelines for the care and use of animals were followed. All procedures performed in studies involving animals were in accordance with the ethical standards of our institution which adhere to the CPCSEA and NIH guidelines.

